## Appendix for "Coordination Between Cytoplasmic and Envelope Densities Shapes Cellular Geometry in *Escherichia coli*"

April 7, 2025

#### Contents

**1 Strains** **1**

**2 Image Processing** **2**

**3 Bayesian Inference** **2**
3.1 Inferring the Total Protein and RNA Mass per Cell . . . . . . . . . . 6
3.2 Inferring Model Parameters . . . . . . . . . . 6

**4 Relating Ribosomal Proteome Allocation $\phi_{rib}$ and the RNA-to-Protein Ratio** **7**

#### 1 Strains

All *Escherichia coli* strains used in this work are described in Table 1. Plasmids with MeshI (GE463) or RelA (GE462) under inducible control were acquired from Addgene and transformed into our wildtype NCM3722 host strain.

| Strain | Source | Description |
|---|---|---|
| NCM3722 | Laboratory of Terence Hwa | WT |
| GE463 | Plasmid from Addgene, AddGeneID:175594 | pMeshI |
| GE462 | Plasmid from Addgene, AddGeneID:175595 | pRelA |

**Table 1:** Strains used in this study.

#### 2 Image Processing

In this work, we directly measured cell size parameters from phase-contrast microscopy images at 100X magnification. We employed an in-house image processing Python pipeline to segment and measure per-cell parameters. The steps of this segmentation algorithm are outlined in Fig. A1. Briefly, individual cells are identified in an image through several rounds of filtering and thresholding. Once cell masks are identified, each is rotated and aligned to a common axis and contours are determined by edge detection, smoothing, and spline interpolation. With a spline interpolation in place, the curvature $k$ along the contour along an $xy$ coordinate system at each point is calculated via

$$k = \frac{x'y'' - y'x''}{(x'^2 + y'^2)^{3/2}} \tag{1}$$

Note that this preserves sign of the curvature. By rotating each cell mask to have the same axis of orientation, we can ensure that the sign of the curvature is consistent between individual cells and images.

With an estimate of the cell curvature in place at each position along the contour, we apply a threshold on this value to identify what portions of the contour correspond to the cell sides ($k \approx 0$) versus the cell caps ($|k| > 0$). We identified the cap regions of the cells as the contour points with radii of curvature greater than 0.5 µm. To compute the cell length dimension, we took the length to be the maximum y coordinate difference between contour points of the two caps. For the width, we computed the average minimum pairwise distance between the two cell side contours. Representative segmented cell peripheries are shown in Fig. A2 for 100 cells from a single wildtype sample grown in a glucose minimal medium. We found that our segmentation protocols yielded measurements that were in line with literature cell size measurements and exhibited the same apparent growth-rate dependence, as shown in Fig. S2(E-H).

#### 3 Bayesian Inference

In this work, we utilize Bayesian statistical methods to systematically propagate all uncertainty from measurements and our model assumptions to generate the final predictions and experimental data presented in this work. In the subsections that follow, we present detailed descriptions of the various components of the inference. In all cases, inferences using literature data and inferences using our suite of experimental measurements were performed independently. Further, we note that *all* inference of model parameters and calculation of estimated membrane protein densities were conducted simultaneously, resulting in a rather large posterior probability distribution that precludes complete enumeration here. However, we invite the reader to examine the full statistical inference models, which are written using the Stan probabilistic programming language [1], and which are available on the paper GitHub repository (`github.com/cremerlab/density_maintenance`). We find that these models, coupled with the inline comments, are easier to parse than a full mathematical statement.

Speaking generally, we sought to inference the probability of a parameter $\theta$ taking on a given value, conditioned on an experimental measurement $y$. This quantity, termed the *posterior probability distribution* $g(\theta \mid y)$ (hereafter called the *posterior*) can be computed using Bayes' rule,

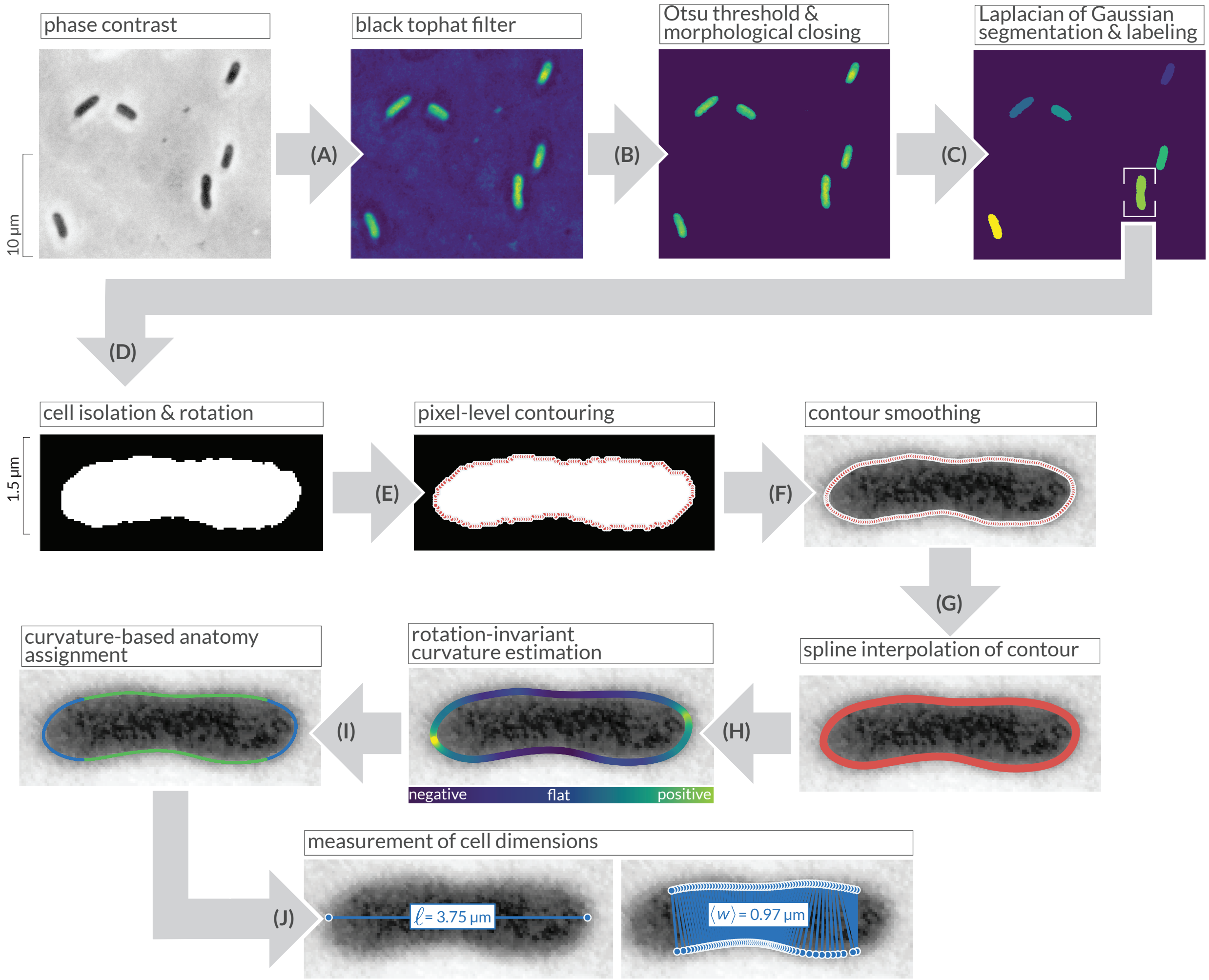


**Figure A1: Segmentation and cell measurement pipeline.** Raw phase-contrast images of cells are passed through (A) a black tophat filter and (B) Otsu thresholding to arrive at emphasized cell boundaries. (C) Laplacian of Gaussian segmentation is then applied to arrive at rough cell masks. For each cell, the sgementation mas is (D) rotated and aligned and the periphery is identified using (E) contouring and (F) Savgol filtering. The isolated contours are then smoothed with (G) spline interpolation. In a clock-wise direction, the (H) curvature of the contour is calculated and used to (I) identify sides and caps. With morphology assigned, cell dimensions are then (J) measured.

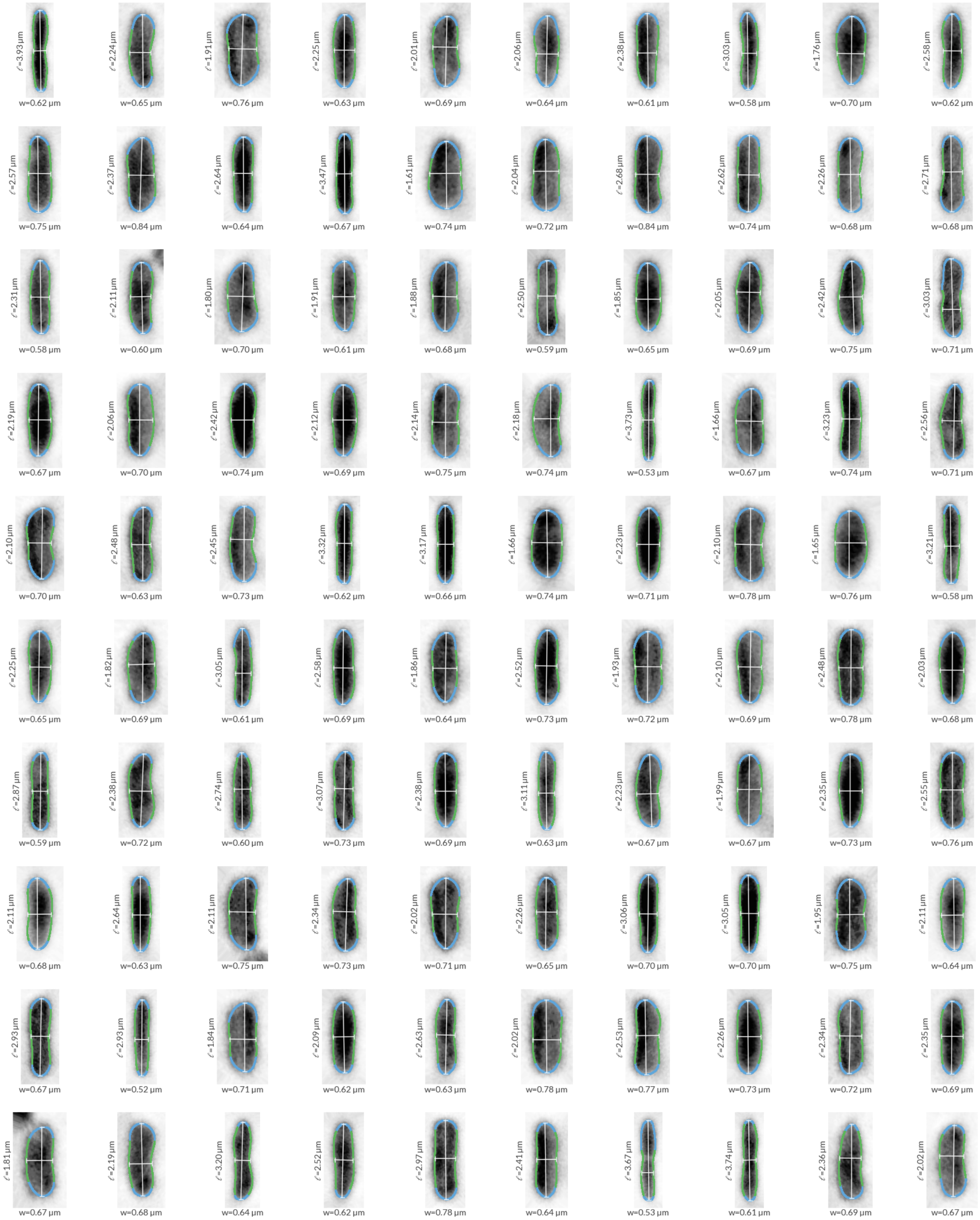


**Figure A2: Example segmentation of *E. coli* cells grown on a glucose minimal medium to steady-state.** Blue and green regions of the contours correspond to the caps and sides of the cells, respectively.

$$g(\theta \,|\, y) = \frac{f(y \,|\, \theta)g(\theta)}{f(y)} \tag{2}$$

where $g$ and $f$ denote probability density functions over parameters and data, respectively. In computing the posterior, one must minimally enumerate the likelihood function $f(y \,|\, \theta)$ and the prior $g(\theta)$. The denominator of Eq. 2 is termed the *evidence* or the *marginalized likelihood* and represents the probability of observing a datum $y$ irrespective of the model. For our purposes, we can treat this as a normalization constant and neglect it. As such, Eq. 2 becomes

$$g(\theta \,|\, y) \propto f(y \,|\, \theta)g(\theta). \tag{3}$$

The likelihood function $f(y \,|\, \theta)$ represents the probability of observing a datum $y$ *given* a particular value of the parameter $\theta$. In all cases for this work, we took the likelihood to have the form of a Gaussian distribution parameterized by a mean $\mu$ and homoscedastic error $\sigma$,

$$f(y \,|\, \theta) = \frac{1}{\sqrt{2\pi\sigma^2}} \exp\left[-\frac{(y-\mu)^2}{2\sigma^2}\right] \Rightarrow f(y \,|\, \theta) \sim \text{Normal}(\mu, \sigma) \tag{4}$$

where we have introduced a short-hand notation. Asserting a Gaussian likelihood makes an assumption about how the measurements are distributed about a mean value $\mu$, but not *what* that mean value is. The remaining subsections of this Appendix section outline how we determine what the mean value of this likelihood function is for a variety of components of our models.

Finally, we must also provide a definition of the prior distribution over the parameter $g(\theta)$. This distribution encapsulates all knowledge we have of what the true parameter value of $\theta$ might be *without* taking the observations into account. This is a critically important point and each prior choice represents the assumptions and domain expertise we employ in crafting these models. In the subsections that follow, we outline the inferential models and provide our prior choices. We used the Probability Distribution Explorer tool (`distribution-explorer.github.io`) to interactively identify parameter values that yielded prior distributions consistent with our domain knowledge.

To compute credible regions of model predictions and fits from the posterior distributions (such as the uncertainty bands in Fig. 3), we employed Markov Chain Monte Carlo (MCMC) sampling techniques implemented through Stan's probabilistic programming framework [1]. For a given model with a Gaussian likelihood as in Eq. 4, we first draw samples of $\mu$ and $\sigma$ from their posterior distributions. Using these posterior parameter samples, we generate posterior predictive datasets $\tilde{y}$ by drawing from the likelihood $f(\tilde{y} \,|\, \theta) \sim \text{Normal}(\mu_{posterior}, \sigma_{posterior})$. We construct credible intervals at the 68% and 95% levels, which approximately correspond to one and two standard deviations in a normal distribution, respectively. This is achieved by finding the appropriate quantiles of the posterior samples for each parameter of interest. The process allows us to construct marginal and joint credible intervals, while simultaneously enabling model validation through posterior predictive checks. By comparing the statistical properties of these simulated datasets $\tilde{y}$ to our observed data $y$, we can diagnose potential model misspecifications, assess the model's predictive performance, and quantify the uncertainty inherent in our parameter estimates.

#### 3.1 Inferring the Total Protein and RNA Mass per Cell

In this work, we combined independent measurements of the total protein and RNA per cell in wildtype *E. coli* with our mass spectrometry measurements to calculate various properties, including the total protein masses in each cellular compartment and the corresponding densities.

We assumed that the total protein per cell $M_{prot}^{(tot)}$ scaled exponentially with the steady-state growth rate $\lambda$,

$$M_{prot}^{(tot)}(\lambda) = \beta_{prot,0} \exp\left(\beta_{prot,1}\lambda\right), \tag{5}$$

where $\beta_{prot,0}$ and $\beta_{prot,1}$ are phenomenological coefficients. Setting this equation as the mean of a Normal likelihood, we inferred the values of these parameters using MCMC setting the following priors:

$$\beta_{prot,0} \sim \text{Inv.Gamma}(\alpha = 13.21, \beta = 1062)\,; \beta_{prot,1} \sim \text{Normal}(0, 1), \tag{6}$$

where Inv.Gamma denotes the Inverse Gamma function. Values for the location and scale parameters of this distribution ($\alpha$ and $\beta$) were chosen such that the upper and lower bounds of the 95$^{\text{th}}$ percentile were 50 fg/cell and 150 fg/cell, respectively.

We also assumed that the total RNA per cell $M_{RNA}^{(tot)}$ scaled exponentially with the steady-state growth rate $\lambda$,

$$M_{RNA}^{(tot)}(\lambda) = \beta_{RNA,0} \exp\left(\beta_{RNA,1}\lambda\right), \tag{7}$$

where $\beta_{RNA,0}$ and $\beta_{RNA,1}$ are the phenomenological coefficients we seek to infer. For this inference, we set the following priors:

$$\beta_{RNA,0} \sim \text{Inv.Gamma}(\alpha = 3.358, \beta = 77.87)\,; \beta_{RNA,1} \sim \text{Normal}(0, 1), \tag{8}$$

where Inv.Gamma denotes the Inverse Gamma function. Values for the location and scale parameters of this distribution ($\alpha$ and $\beta$) were chosen such that the upper and lower bounds of the 95$^{\text{th}}$ percentile were 10 fg/cell and 100 fg/cell, respectively.

#### 3.2 Inferring Model Parameters

As derived in the main text, the central prediction of the density maintenance model is that the cellular surface-to-volume ratio $S_A/V$ is related to the cellular composition and proteome partitioning between compartments via

$$S_A/V = \frac{\kappa \psi_{mem}}{2\left[1 - \psi_{mem} - \psi_{peri} + \beta\phi_{rib}\right]} \tag{9}$$

where $\psi_{mem}$ is the proteome partitioning towards the membrane, $\psi_{peri}$ is the partitioning toward the periplasm, $\kappa$ is the cytoplasm-membrane density ratio, and $\phi_{rib}$ is the ribosomal proteome allocation, related to the RNA-to-protein ratio via the proportionality constant $\beta$, derived in the next section. To test our model, we sought to estimate the value of $\kappa$ that could describe how the $S_A/V$ scaled as $\phi_{rib}$ was varied, requiring us to also estimate how $\psi_{mem}$ and $\psi_{peri}$ depended on $\phi_{rib}$.

There are many different functional forms one could use to describe how the $\psi$'s depended on $\phi_{rib}$. We made the empirical assumption that the membrane proteome partitioning $\psi_{mem}$ was linearly dependent on

$\phi_{rib}$ with the form

$$\psi_{mem}(\phi_{rib}) = \beta_{0,\psi_{mem}} + \beta_{1,\psi_{mem}}\phi_{rib} \tag{10}$$

where $\beta_{0,\psi_{mem}}$ and $\beta_{1,\psi_{mem}}$ are phenomenological constants we sought to infer. For this inference, we defined the priors to be

$$\beta_{0,\psi_{mem}} \sim \text{Beta}(\alpha = 1.262, \beta = 5.967)\,;\, \beta_{\psi_{mem}} \sim \text{Normal}(0, 1), \tag{11}$$

where Beta denotes the Beta distribution, which is defined on the interval [0, 1]. Values for the location $\alpha$ and scale $\beta$ parameters of the Beta distribution were chosen such that the lower and upper bounds of the 95$^{\text{th}}$ percentile were 0.01 and 0.5, respectively.

For the periplasmic partitioning, we empirically modeled an exponential dependence on $\phi_{rib}$, having the form

$$\psi_{peri}(\phi_{rib}) = \beta_{0,\psi_{peri}} \exp\left(\beta_{1,\psi_{peri}}\phi_{rib}\right), \tag{12}$$

where the priors for the phenomenological constants $\beta_{0,\psi_{peri}}$ and $\beta_{1,\psi_{peri}}$ were similarly defined to be

$$\beta_{0,\psi_{peri}} \sim \text{Beta}(\alpha = 1.262, \beta = 5.967)\,;\, \beta_{1,\psi_{peri}} \sim \text{Normal}(0, 10), \tag{13}$$

where a broader prior on $\beta_{1,\psi_{peri}}$ was defined to permit a stronger dependence on $\phi_{rib}$. Bounds for $\alpha$ and $\beta$ in the Beta distribution were empirically chosen such that the lower and upper bounds of the 95$^{\text{th}}$ percentile were 0.01 and 0.5, respectively

These dependencies were inferred simultaneously with our inference of $\kappa$, given our direct measurement of $S_A/V$ and $\phi_{rib}$ given Eq. 9. For this inference, we assigned a prior distribution over $\kappa$ to be

$$\kappa \sim \text{Inv.Gamma}(\alpha = 2.863, \beta = 140.1), \tag{14}$$

where the values for the location $\alpha$ and scale $\beta$ parameters were chosen such that the lower and upper bounds of the 95$^{\text{th}}$ percentile were 20 µm$^{-1}$ and 250 µm$^{-1}$, respectively.

### 4 Relating Ribosomal Proteome Allocation $\phi_{rib}$ and the RNA-to-Protein Ratio

In the main text, we make the assertion that the ribosomal proteome fraction $\phi_{rib}$ is related to the RNA-to-protein ratio $M_{RNA}^{(tot)}/M_{prot}^{(tot)}$ via the proportionality constant $\beta$. This constant can be estimated directly [2,3] given our knowledge of the RNA and protein content of the ribosome [2,4].

The vast majority of total cellular RNA is ribosomal [2], allowing us to state

$$M_{rRNA}^{(tot)} \approx \alpha M_{RNA}^{(tot)}, \tag{15}$$

where $M_{rRNA}^{(tot)}$ is the total rRNA mass of the cell, $M_{RNA}^{(tot)}$ is the total RNA mass and $\alpha$ is a fractional quantity between 0 and 1. We can make the assumption that approximately all rRNA is associated with ribosomal particles, allowing us to calculate this mass $M_{rRNA}$ as

$$M_{rRNA}^{(tot)} = m_{rRNA}N_{rib}, \tag{16}$$

where $m_{rRNA}$ is the total mass of rRNA associated with a single ribosome and $N_{rib}$ is the total number of ribosomes per cell. Assuming that all ribosomal proteins are within assembled ribosomes, the total number of ribosomes $N_{rib}$ can be calculated from the ribosomal allocation $\phi_{rib}$ and the total protein mass $M_{prot}^{(tot)}$ as

$$N_{rib} = \frac{\phi_{rib} M_{prot}^{(tot)}}{m_{rib}}, \tag{17}$$

where $m_{rib}$ is the total proteinaceous mass of a single ribosome.

We can combine Eq. 15 - Eq. 17 and solve for the total RNA-to-protein ratio $M_{RNA}^{(tot)}/M_{prot}^{(tot)}$ to yield

$$\frac{M_{RNA}^{(tot)}}{M_{prot}^{(tot)}} = \frac{m_{rRNA}}{\alpha m_{rib}} \phi_{rib} = \beta \phi_{rib}, \tag{18}$$

where $\beta$ is the proportionality constant relating ribosomal allocation and the RNA-to-protein ratio. Using our knowledge of the total length of rRNA ($\approx$ 4500 nt) and total size of ribosomal proteins ($\approx$ 7500 AA [3]), the precise value of $\beta$ can be calculated as

$$\beta = \overbrace{\left[\frac{4500\,\text{RNA nt}}{\text{ribosome}} \cdot \frac{340\,\text{Da}}{\text{RNA nt}}\right]}^{m_{rRNA}} \cdot \underbrace{\left[\frac{\text{ribosome}}{7500\,\text{AA}} \cdot \frac{\text{AA}}{110\,\text{Da}}\right]}_{m_{rib}} \cdot \underbrace{\left[\frac{1}{0.85}\right]}_{\alpha} \approx 2.18, \tag{19}$$

which is in close alignment with other calculations [2, 3].
